# Computed cancer interactome explains the effects of somatic mutations in cancers

**DOI:** 10.1101/2022.01.21.477304

**Authors:** Jing Zhang, Jimin Pei, Jesse Durham, Tasia Bos, Qian Cong

## Abstract

Protein-protein interactions (PPIs) are involved in almost all essential cellular processes. Perturbation of PPI networks plays critical roles in tumorigenesis, cancer progression and metastasis. While numerous high-throughput experiments have produced a vast amount of data for PPIs, these datasets suffer from high false positive rates and exhibit a high degree of discrepancy. Coevolution of amino acid positions between protein pairs has proven to be useful in identifying interacting proteins and providing structural details of the interaction interfaces with the help of deep learning methods like AlphaFold (AF). In this study, we applied AF to investigate the cancer protein-protein interactome. We predicted 1,798 PPIs for cancer driver proteins involved in diverse cellular processes such as transcription regulation, signal transduction, DNA repair and cell cycle. We modeled the spatial structure for the predicted binary protein complexes, 1,087 of which lacked previous 3D structure information. Our predictions offer novel structural insight into many cancer-related processes such as the MAP kinase cascade and Fanconi anemia pathway. We further investigated the cancer mutation landscape by mapping somatic missense mutations (SMMs) in cancer to the predicted PPI interfaces and performing enrichment and depletion analyses. Interfaces enriched or depleted with SMMs exhibit different preferences for functional categories. Interfaces enriched in mutations tend to function in pathways that are deregulated in cancers and they may help explain the molecular mechanisms of cancers in patients; interfaces lacking mutations appear to be essential for the survival of cancer cells and thus may be future targets for PPI modulating drugs.

## Introduction

Cancer development and progression are characterized by the combined deregulation of multiple cellular processes resulting in unrestricted proliferation and replication potential, insensitivity and resistance to anti-growth signals and apoptosis, promotion of angiogenesis, and the ability to invade and metastasize [1]. Protein–protein interactions (PPIs) are important in cancer biology as they are the foundations for correct formation of functional protein complexes and serve to relay cell growth and death signals between components in cancer-related signaling pathways [2]. PPIs are traditionally regarded as high-hanging fruits for drug development [3], but cancer-enabling PPIs are pursued as promising targets of developing drugs that aim to disrupt their interfaces in recent years [4]. For example, PPI modulators for PD-1/PD-L1 and Bcl-2/Bax are approved for cancer treatment, and even more candidates are in the clinical trials [5]. Large-scale high-throughput experimental studies utilizing techniques such as affinity purification and chemical cross-linking have been used to detect PPIs in human and other organisms at the whole proteome level [6-8] or for cancer-related proteins [9, 10]. However, high-throughput PPI datasets often have high false positive and false negative rates and show considerable inconsistencies between studies [11, 12].

Cancer genomics studies aim to identify genetic differences between cancer cells and normal cells by next-generation sequencing techniques [13]. Such studies have advanced our understanding of the molecular mechanisms of cancer growth, metastasis, and drug resistance in various cancer types and expanded the landscape of cancer-related genes and their genetic alterations [14-18]. Genome sequencing is also used to detect known disease-causing genetic variations that provide valuable information for making clinical decisions for cancer patients [19]. Cancer-causing genetic alterations lead to activation of oncogenes or deactivation of tumor suppressor genes, which can occur through structural changes of chromosomes, fusion of individual genes, as well as germline or somatic mutations at individual nucleotide positions [20].

Somatic mutations in cancer have been continuously discovered in large-scale sequencing of genomes and exomes [14]. The landscape of somatic mutations in cancer provides a useful resource to identify recurrently mutated cancer driver genes in various cancer types [15, 21, 22]. Somatic mutations providing a selective growth advantage are classified as cancer driver mutations, including a small fraction of mutations with high occurrence frequencies (mutation hotspots) [23]. However, the majority of somatic mutations occur at low frequencies in cancers and belong to the category of unknown clinical significance. It remains a challenge to interpret their functional significance and to differentiate cancer driver mutations from the passenger somatic mutations. Mapping mutations on three dimensional structures can assist our interpretation of these mutations [24, 25]. A significant fraction of somatic mutations occurs at the interface of PPIs [26, 27]. As PPIs play crucial roles in diverse cellular processes, these somatic mutations could have significant functional and clinical consequences, e.g., by disrupting the correct formation of protein complexes and interrupting their function [19], or by improperly increasing the affinity between proteins and leading to deregulation of signaling pathways [28-30]. Therefore, accurate prediction of PPI interfaces could help interpret the clinical significance of somatic mutations and provide targets for cancer drug development.

Recent breakthroughs in deep learning (DL) methods for protein structure predictions have significantly expanded the capability of obtaining high-resolution protein structures [31] and interactions [32]. We recently found that the DL methods developed for monomeric protein structure prediction, i.e. RoseTTAFold [33] and AlphaFold (AF) [34], can be used to identify interacting protein pairs with an accuracy better than large-scale experimental studies in yeast [35]. Application of these DL methods to candidate PPIs identified in large-scale studies not only allows us to identify a confident set of PPIs from noisy high-throughput experimental data, but also provides high-quality 3D structure models for the protein complexes. In this study, we applied AF to investigate the PPI network involving cancer driver proteins, a step towards building a structurally resolved cancer interactome (Fig. S1). We integrated the interactome with somatic missense mutations (SMMs) in cancer cells from 12,711 patients [36], and illustrated that the structural information of cancer interactome can help determine potential cancer driving mutations, explain their mechanistic consequences to protein complex function and signaling transduction, and provide targets for cancer treatment.

## Results and Discussions

### The protein-protein interaction network around cancer-driving proteins

Recent cancer genomic studies identified 723 cancer-driving proteins that contain mutations causally implicated in cancer according to the Catalogue of Somatic Mutations in Cancer (COSMIC) Cancer Gene Census (CGC) database (Oct 5th, 2021) [37]. These cancer-driving proteins can be classified as the products of oncogenes, tumor suppressor genes (TSGs) and genes that contribute to cancer progression by fusion with other genes. We excluded the cancer-driver genes functioning solely by gene fusion from this study because these genes mainly contribute to cancer through transcriptional regulations and not through mutations in their protein products that can disrupt their interaction with other molecules.

We extracted the putative interacting human proteins for the remaining 587 CGC cancer driver proteins (Table S1) from various PPI databases [8, 38-40]. Out of 705,087 putative human PPIs from these databases, 103,756 involve cancer driver proteins. Cancer driver proteins tend to have significantly (P-value: 2.54e-72) more putative interacting partners than other human proteins (Fig. S2), consistent with the well-known fact that many cancer-drivers are hubs of PPIs [9, 41, 42]. Since most of these PPIs were identified from large-scale experiments that are prone to false positives [11, 12], we extracted 137,222 PPIs supported by multiple high-throughput experiments or at least one low-throughput study (see Methods). Among them, we focused on 26,947 PPIs involving at least one cancer driver protein (Table S2). We used AF to model the spatial structures for 24,152 of these candidate PPIs, and the remaining 2,795 (10%) were excluded from this study due to difficulty in obtaining multiple sequence alignments (MSAs) or limitation in GPU memory (details in methods).

We previously showed that although AF was developed to model monomeric proteins, it can be extended to accurately model 3D structures of protein complexes [35]. In addition, the top AF contact probability between residues of a protein pair can be used to distinguish true PPIs from false ones in yeast [35]. This is still true for the set of cancer-related human PPIs. We computed the AF contact probabilities for 8,000 candidate PPIs involving cancer drivers and for 8,000 pairs of cancer drivers and random proteins (Fig. 1A). Among the pairs with AF contact probability above 0.9, candidate PPIs are enriched by 18-fold compared to random pairs, and at a contact probability cutoff of 0.8, candidate PPIs are enriched by 5.5-fold. Among the 24,152 binary protein complexes we modeled, 1,459 have closely-related experimental structures, allowing us to estimate the accuracy of the predicted interface. The vast majority (96.4% for contact probability ≥ 0.9 and 92.6% for contact probability ≥ 0.8) of the residue pairs showing high contact probability are less than 12Å away in experimental structures (Fig. 1B). When the predicted contacts are evaluated separately for each protein complex, the majority (≥ 60%) of the high-confidence (probability ≥ 0.8) inter-protein contacts agree with the experimental structure for 93% AF complex models (Fig. 1C).

**Figure 1.**
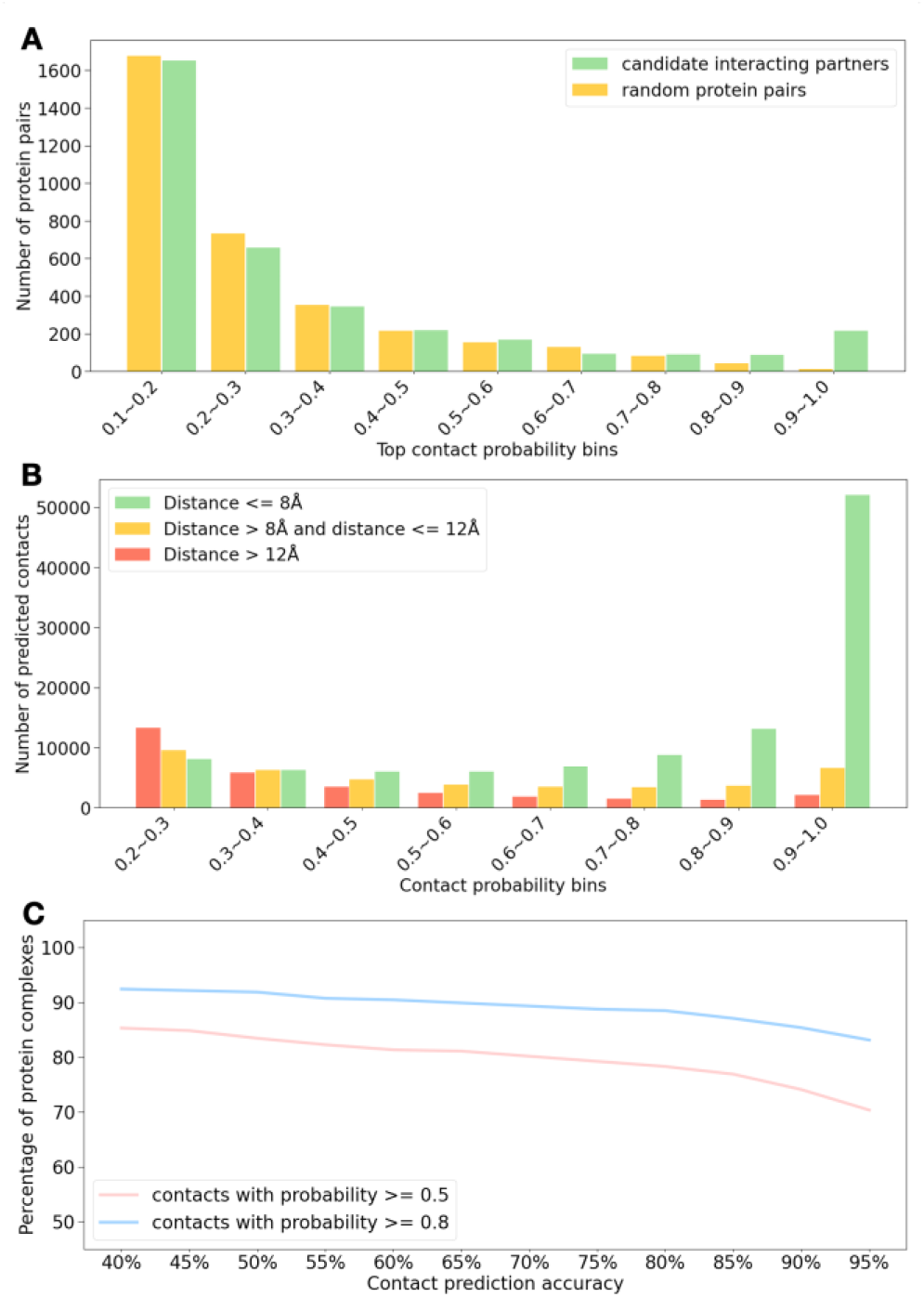
Benchmark for the performance of complex modeling by AlphaFold. **(A)** AF top contact probability can distinguish interacting proteins from random pairs. 8,000 pairs of cancer driver and its candidate interacting partner and 8,000 pairs of cancer driver and a random protein were modeled by AF. These pairs are partitioned to bins by the top contact probability, and the number of pairs in each bin is shown. **(B)** Distribution of residue-residue distances in closely related experimental structures for AF-predicted contacts in 1,459 protein pairs with close homologous structures in PDB. The predicted contacts are binned according to the AF contact probability. **(C)** Distribution of contact prediction accuracy for 1,459 AF models with close templates in PDB. The contact prediction accuracy for an AF model was calculated as the percentage of correctly predicted contacts and a predicted contact was considered correct if the two residues forming this contact were less than 12Å in closely-related experimental structures.

These benchmarks suggest that AF contact probability can be used as a measure to select a set of confident PPIs involving cancer drivers among the large-scale experimental data that are prone to errors, and it can model the binary complexes between these proteins with high accuracy. In this study, we focused on 1,798 (Table S3) cancer-related PPIs with top AF contact probability above 0.8, and we visualized the network formed by these PPIs with Cytoscape [43]. These protein complexes are deposited to ModelArchive under accession ???. Since this PPI network centers around cancer drivers and many of these interactions may be related to oncogenesis or cancer progression, we term this network “cancer interactome” [44]. We performed clustering analysis [45] of the “cancer interactome” to partition the network into groups of functionally related proteins that are linked by direct physical interactions. The resulting clusters (Tables S4-S34), excluding small clusters with less than 10 proteins, are shown in Fig. 2, where proteins are represented as dots and interacting proteins are connected by edges.

**Figure 2.**
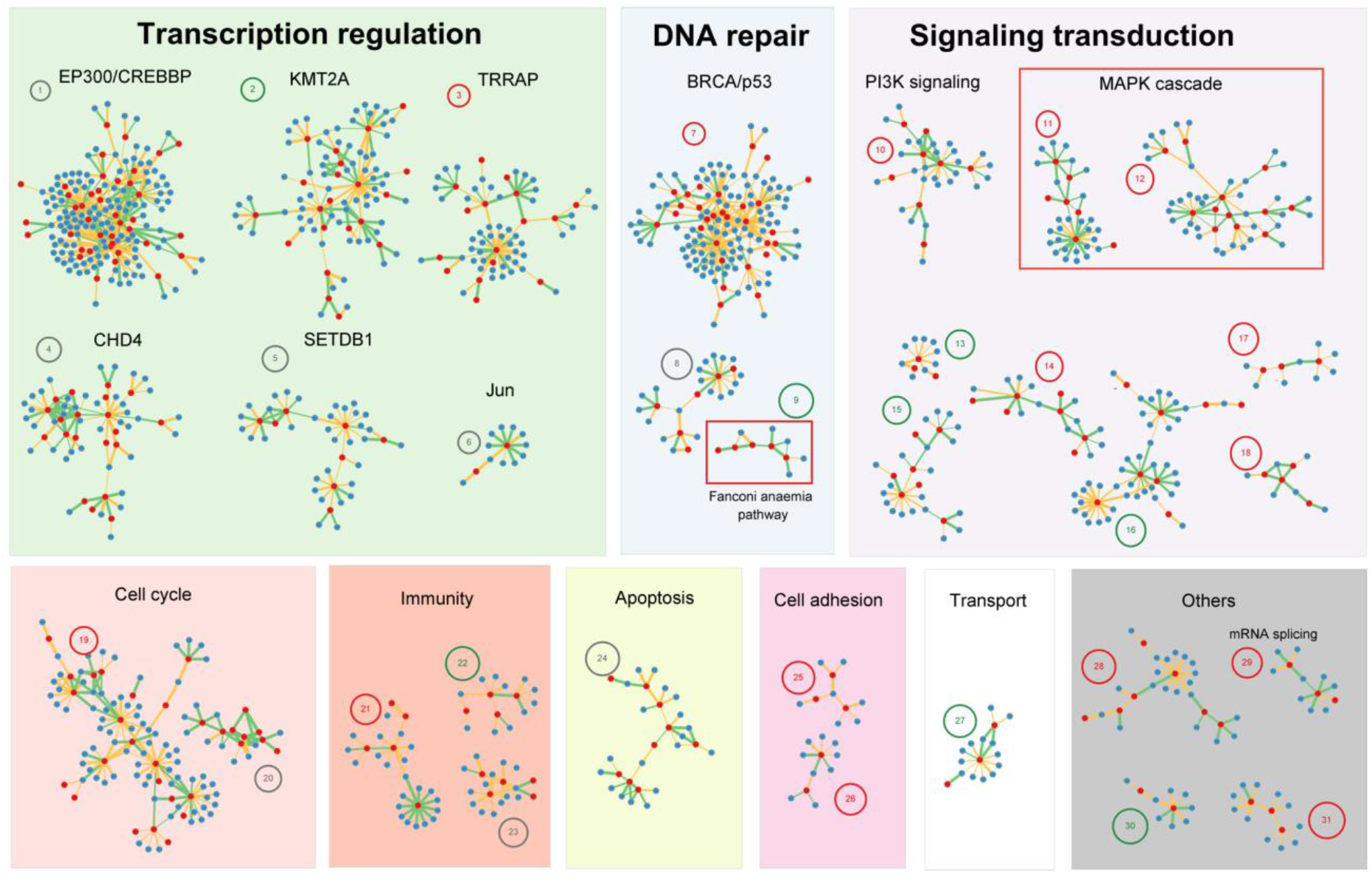
Clusters of proteins in the cancer interactome. Dots represent proteins and edges connect proteins that are predicted to interact (AF contact probability ≥ 0.8). Clusters are labeled by its predominant functional category or the hub protein which forms the largest number of interactions within the cluster. Cancer driver proteins are colored red and their interacting partners are colored blue. Edges representing predicted interactions without experimentally determined 3D structures are colored orange and interactions with homologous templates in PDB are colored green. Edge width is positively correlated with AF contact probability. A number inside a circle is used to label each cluster, and information about proteins in each cluster is in supplemental Tables S4-S34. We color this label red or green if cancer drivers in this cluster are enriched in or depleted of SMMs in cancer, respectively. Pathways that are discussed in details, i.e. MAPK cascade and FA pathway, are placed in red boxes.

The most predominant clusters involve transcriptional regulators such as EP300 and CREBBP that regulate the expression of a wide range of genes through chromatin remodeling [46], signaling transduction such as the PI3K and the MAPK pathways, and the DNA repair pathways involving proteins like BRCA1 and BRCA2. Other processes involved in the cancer interactome include cell cycle, immunity, apoptosis, cell adhesion and transport. Although all of these processes are known to play vital roles in cancer [1], our study provides novel structural information for a significant fraction of these interactions. Only 711 out of the 1,798 confident complex models (39.5%) have homologous complex structures (BLAST e-value < 0.00001, green edges in Fig. 2) that were determined experimentally according to PDB. The remainder (60.5%) (orange edges in Fig. 2) are novel structures that could provide insights into the functional mechanisms of these proteins.

### Somatic mutational landscape of the cancer interactome

We obtained SMMs in 12,711 cancer patients from the OncoVar database [36]. We assigned these SMMs to 17,515 reviewed human entries in UniProt that can be unambiguously and perfectly mapped to the reference genome GRCh38, and the average SMM rate for these proteins is 0.00146% per position per patient. The SMM rate in cancer driver proteins is 0.00181%, significantly (P-value < 1e-320) higher than other genes, since many of these proteins harbor mutation hotspots.

However, not all the cancer drivers are highly mutated in this pan-cancer dataset. We analyzed the SMM rate of proteins from each functional cluster in our cancer interactome (Fig. 2). While some pathways (labeled by red circles around the numbers), such as PI3K and MAPK pathways, are highly enriched in cancer SMMs, some are significantly depleted (labeled by green circles around the numbers) of such SMMs (Table S35). The seemingly paradox of SMM depletion of some cancer driver proteins can be explained by the complexity of oncogenesis, cancer progression, and cancer cell evolution during treatment. For example, the well-known Breast cancer type 2 susceptibility protein, BRCA2, indeed has a relatively low SMM rate both in the pan-cancer dataset (0.00133%) and the breast cancer subset (0.00075%), consistent with previous findings [47]. Instead, most studied cancer-causing mutations of BRCA2 are germline (inherited) variations that can increase the chance for their carriers to develop cancer [48]. The same is true for proteins in the Fanconi anemia (FA) pathway involved in DNA repair, which also have low SMM rates, and genetic variants in them can predispose the carriers to higher risk of oncogenesis. DNA repair is an important process for cancer cells, especially for them to survive during radiotherapy and chemotherapy that kills cancer cells mostly by damaging their DNA [49]. Therefore, although mutations in them may contribute to carcinogenesis by increasing the rate of mutations in the genome, accumulating excessive SMMs in DNA repair proteins is likely not beneficial for most cancer types.

The PPI interfaces in our cancer interactome contain a total of 46,472 amino acid residues. 11,726 cancer SMMs mapped to these interface residues, indicating a SMM rate of 0.00199%, significantly (P-value = 1.2e-17) higher than the overall SMM rate for cancer drivers. SMMs on a PPI interface may disrupt or enhance the interaction between two proteins [4, 24]. While a considerable portion of SMMs might be passenger mutations, the enrichment or depletion of SMMs in a PPI interface could imply a role of this PPI in cancer development. Enrichment of SMMs in a PPI interface suggests that disturbing this PPI could facilitate oncogenesis and cancer progression [25, 50], and depletion of SMMs in a PPI interface indicates that maintaining this interaction is likely beneficial to cancer cells.

We identified 176 PPI interfaces (Table S36) that are significantly enriched in SMMs, and 65 interfaces that are significantly depleted (Table S37) of SMMs. We performed GO term enrichment analysis to reveal the functional difference of proteins with these two types of interfaces (Fig. 3A, Table S38, S39). The overrepresented biological processes by proteins with SMM-rich interfaces but not by proteins with SMM-depleted interfaces include signal transduction that controls cell proliferation and growth as represented by MAPK and PI3K cascades. Interacting proteins involved in cell differentiation or organ development also tend to accumulate SMMs in their interfaces. An interesting overrepresented functional category by SMM-rich PPI is antigen processing and presentation, a process that, if disturbed, may contribute to immune evasion by introducing SMMs to key interfaces.

**Figure 3.**
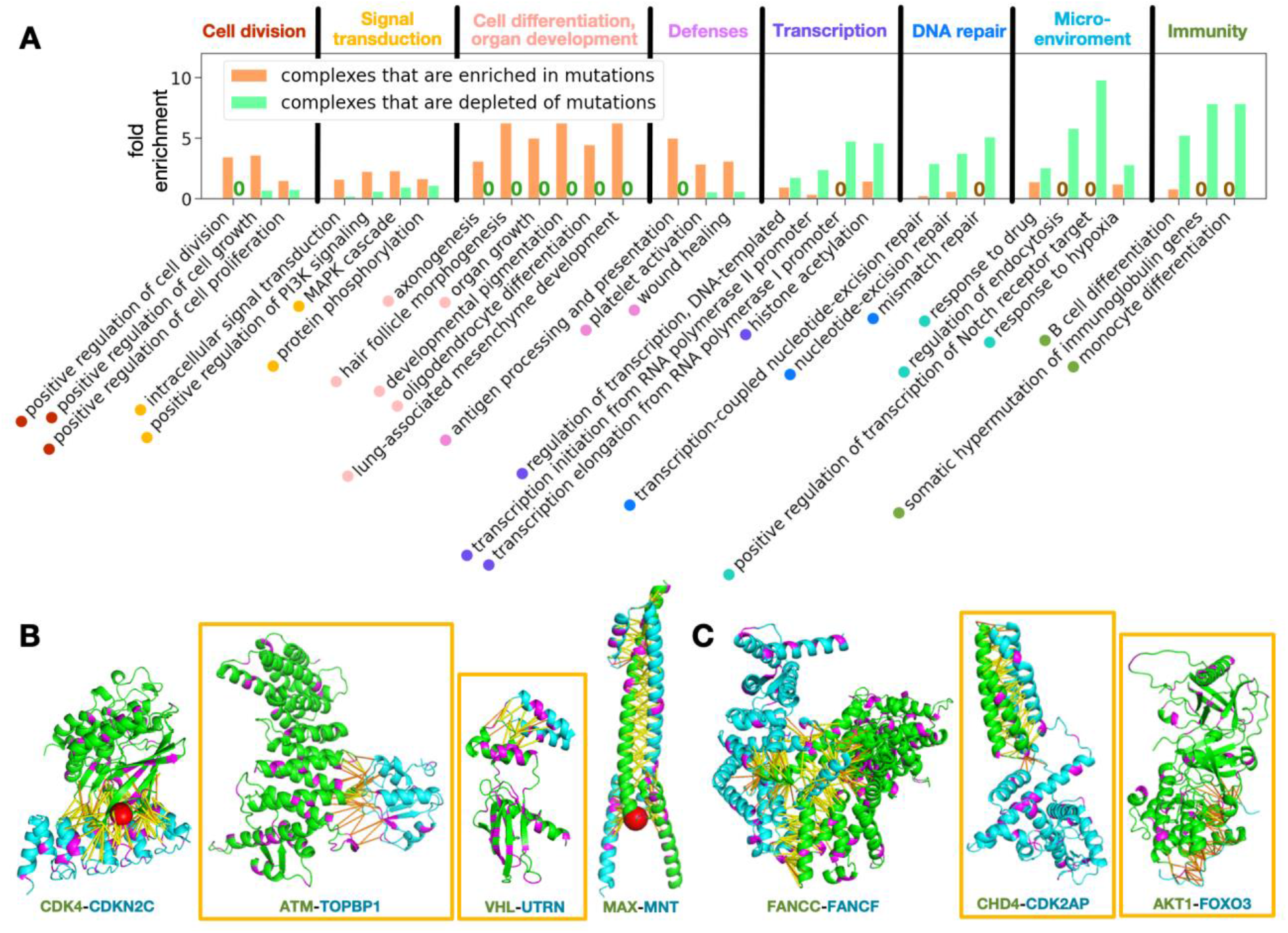
Protein-protein interaction interfaces that are enriched in or depleted of somatic missense mutations in cancer. **(A)** Gene Ontology (GO) terms that are significantly overrepresented by proteins with SMM-rich or SMM-depleted interfaces. For each GO term, we show its fold enrichment among two sets of proteins as bar plots: (1) proteins with SMM-rich interfaces (SMM-rich set, orange bars) and (2) proteins with SMM-depleted interfaces (SMM-depleted set, green bars). Fold enrichment is defined as the fraction of proteins associated with this GO term in the target set (either SMM-rich set of SMM-depleted set) divided by the fraction of proteins with this GO term among all proteins in our cancer interactome. Two bars are expected for each GO term: one orange bar for its fold enrichment in the SMM-rich set and one green bar for its fold enrichment in the SMM-depleted set. However, a bar can be missing if the fold enrichment of a GO term in one set is 0, and we marked such cases as “0” above the x-axis instead of a bar. Annotation of the GO terms are labeled below the bar plot and they are manually partitioned to larger functional groups (separated by black lines) as indicated by the color of the dots before labels and the texts above the bar plot. **(B)** Examples of PPI interfaces that are enriched in SMMs. **(C)** Examples of PPI interfaces that are depleted of SMMs. For both **(B)** and **(C)**, positions with SMMs are colored in magenta, and mutation hotspots are shown as red balls. Predicted contacts are shown in sticks (yellow: probability score ≥ 0.9; orange: 0.5 ≤ probability score < 0.9). Names of the proteins are labeled below the complex structure. Protein pairs without homologous complex structures in PDB are placed inside orange boxes.

In contrast, a different set of functions are overrepresented by complexes with SMM-depleted interfaces (groups on the right of Fig. 3A), including DNA transcription, DNA repair, and responses to drugs and hypoxia. All these processes are likely important for the survival and proliferation of cancer cells: SMMs in these interfaces are expected to have a negative impact on the fitness of cancer cells and are thus selected against in cancer cell evolution. Intriguingly, proteins involved in B-cell and monocyte differentiation also tend to avoid SMMs in their PPI interfaces. Both B-cell and monocyte have complicated roles in cancer, and they can display both pro-tumor and anti-tumor functions depending on a number of factors such as the subtype of B-cell and monocyte, and the microenvironmental cues [51, 52]. Our result is more consistent with a pro-tumor effect of B-cell and monocyte, and cancer cells possibly could evade the anti-tumor effects of these immune cells through other strategies.

Integration of our structurally resolved cancer interactome and the cancer SMM landscape can help explain how SMMs contribute to oncogenesis and cancer progression by affecting the interactions between proteins and the function of protein complexes. A deeper understanding of cancer SMMs may aid in the design of precision therapies. In addition, interfaces that are depleted of SMMs likely play a vital role for the survival of cancer cells, and some of them can be promising drug targets for cancer treatment if disruption of these interfaces do not drastically affect the survival of normal cells. In the following sections, we illustrate these points with carefully studied examples.

### Protein-protein interfaces enriched in mutations reveal mechanisms of cancer

One leading cause of SMM enrichment in a PPI interface is the presence of a mutation hotspot in the interface. For example, the oncogene kinase CDK4 has a hotspot residue R24 [53] residing in the interface between CDK4 and the INK4 family of inhibitors such as CDKN2C (a tumor suppressor) [54, 55]. SMMs involving R24 of CDK4 may abolish the interaction between CDK4 and its inhibitor [56], resulting in constitutive kinase activity (Fig. 3B, left) that promotes the progression of cell cycle and leads to transition from G1 to S phase.

More interestingly, our models allowed us to identify interfaces where multiple cancer SMMs clustered: although SMMs do not occur to each residue as frequently as mutation hotspots, they collectively highlight the potential importance of this interface. One example is the interaction between the kinase ATM and one of its substrates, TOPBP1 (Fig. 3B, middle left), both of which are multidomain proteins functioning in DNA repair through homologous recombination [57, 58]. AF predicted the interaction between them to be mediated by the N-terminal α-helical repeat domain of ATM (residues 1-400) and the fourth BRCT domain of TOPBP1 (residues 340-451). Enrichment of SMMs in this interface is highlighted by a loop region in ATM (residues 290-303) that harbors seven residues with cancer SMMs. One of the SMMs P292L has been found to be a likely pathogenic mutation associated with lymphoma [59, 60], suggesting the importance of this interface in cancer.

The interface between utrophin (UTRN) and the tumor suppressor VHL is also enriched with SMMs but without hotspot residues. UTRN mutations have been found in multiple cancers [61], and *UTRN* has been suggested to be a potential TSG as its decreased expression was associated with various cancers including lung cancer, colon cancer, and melanoma [61-63]. The predicted interaction interface between UTRN and VHL involves an α-helical hairpin of UTRN (residues 2841-2876) and the C-terminal helical domain of VHL (residues: 156-213). The interaction between VHL and UTRN was identified in high-throughput experiments and detailed experimental characterization of the UTRN-VHL interaction is still lacking. Our model of the UTRN-VHL complex (Fig. 3B, middle right) may facilitate further experimental studies to validate this interaction and probe its potential role in cancer.

The effect of SMMs at the PPI interfaces may be challenging to interpret even in the presence of high-quality structure models. One reason is that a protein can interact with multiple partners through the same interface. For example, the transcription factor MAX has a mutation hotspot residue R60 in its interface with multiple partners such as MYC, MNT, MXI1, and MXD1. Our models of MAX and its binding partners, in agreement with experimental structures, show that the protein pairs dimerize through their leucine zipper domains (Fig. 3B, right). *MYC* is a well characterized oncogene as the MYC-MAX complex activates the expression of genes to promote cell proliferation [64]. However, *MAX* has been classified as a tumor suppressor as it was found recurrently inactivated in several types of cancers [65-67]. It was proposed that the tumor suppressor function of MAX is not related to its interaction with MYC, but is dependent on the transcriptional repression complexes formed between MAX and other transcription factors such as MXD, MNT and MGA [68]. Thus, the functional impact of SMMs in MAX including those at the hotspot R60 should be interpreted with caution as MAX is involved in complexes with different roles in cancer. Cancer-causing SMMs in MAX likely exert their effect by disrupting the MAX complexes functioning as transcriptional repressors and not by disrupting the MYC-MAX complex.

### Protein-protein interfaces depleted in somatic mutations may serve as drug targets

As shown in our function enrichment analysis, the interface between proteins in DNA repair pathways are often depleted of SMMs in cancers. One such example is the FANCC-FANCF complex (Fig. 3C, left), two subunits of the FA core complex. While these two proteins bind through a large interface, SMM rate in the interface is only half of the average.

Another example of SMM-depleted interface is between CHD4 and CDK2AP1, core components of the nucleosome remodeling and deacetylase complex that plays an important role in epigenetic transcriptional repression during double-strand break DNA damage repair [69, 70]. CHD4 is classified as an oncogene product as its recruitment to the damage sites in hypermethylated CpG island promoter regions helps maintain the silencing of tumor suppressor genes [70]. A recent study using cross-linking mass spectrometry (CLMS) mapped the interaction site between CHD4 and CDK2AP1 to the C-terminal region of CHD4 [71]. AF prediction of the CHD4-CDK2AP1 interface is consistent with the CLMS experiments and yet provides structural details of this interaction, where both CHD4 and CDK2AP1 contribute an α-helix hairpin to form a 4-helical bundle domain (Fig. 3C, middle). Depletion of SMMs in this interface suggests that the CHD4-CDK2AP1 interaction is crucial to maintain the oncogenic role of CHD4.

We also found SMM-depletion in the AKT1-FOXO3 interface (Fig. 3C, right). The Ser/Thr kinase AKT1, encoded by an oncogene, promotes cell growth and proliferation [72, 73]. FOXO3, in contrast, is a transcription factor acting as a tumor suppressor [74]. AKT1 phosphorylation of FOXO3 promotes the transport and sequestration of FOXO3 in cytoplasm and eventual degradation [75]. The AKT1-FOXO3 interaction interface is mapped to the N-terminal segment (residues 14-36) of FOXO3 that does not have a single SMM in our pan-cancer datasets. Although there are a couple SMM in AKT1 mapping to its interface with FOXO3, not a single mutation hotspot locates in this interface. The depletion of SMMs at this interface suggests that the negative regulation of tumor suppressor FOXO3 through AKT1 is important for cancer cell survival, and drugs to disrupt this interface could offer a therapeutic opportunity in cancer treatment.

### Protein-protein interactions in the MAP kinase pathway

MAP kinases are key components of evolutionarily conserved signaling networks involved in diverse cellular processes such as cell growth, proliferation and apoptosis [76]. The MAPK signaling networks involve sequential phosphorylation steps: MAPK kinase kinases (MAP3Ks, MEK kinases, or MKKKs) phosphorylate and activate MAPK kinases (MAP2Ks, MEK, or MKKs), MAPK kinases phosphorylate and activate MAPKs, and MAPKs phosphorylate downstream kinases and other substrates. Human genome encodes ∼20 MAP3Ks, 7 MAP2Ks, and 11 MAPKs that are part of the integrated signaling networks to respond to diverse stimuli and to control critical functions in virtually all cells.

Several members in the MAPK signaling networks are classified as cancer drivers, including the MAP kinase MAPK1, three MAP2Ks (MAP2K1, MAP2K2 and MAP2K4), and six MAP3Ks (MAP3K1, MAP3K1, MAP3K13, BRAF, ARAF, and RAF1). Gain-of-function SMMs in oncogene products such as BRAF and RAS [77, 78] are frequently observed in certain cancers as they promote cell growth through the RAS/RAF/ERK pathway. Our predictions capture many interactions between components of the consecutive steps in the MAPK cascades (Fig. 4). We also observed MAPK1’s involvement in many different complexes, suggesting that MAPK1 is a hub protein with a diverse range of substrates and other binding partners.

**Figure 4.**
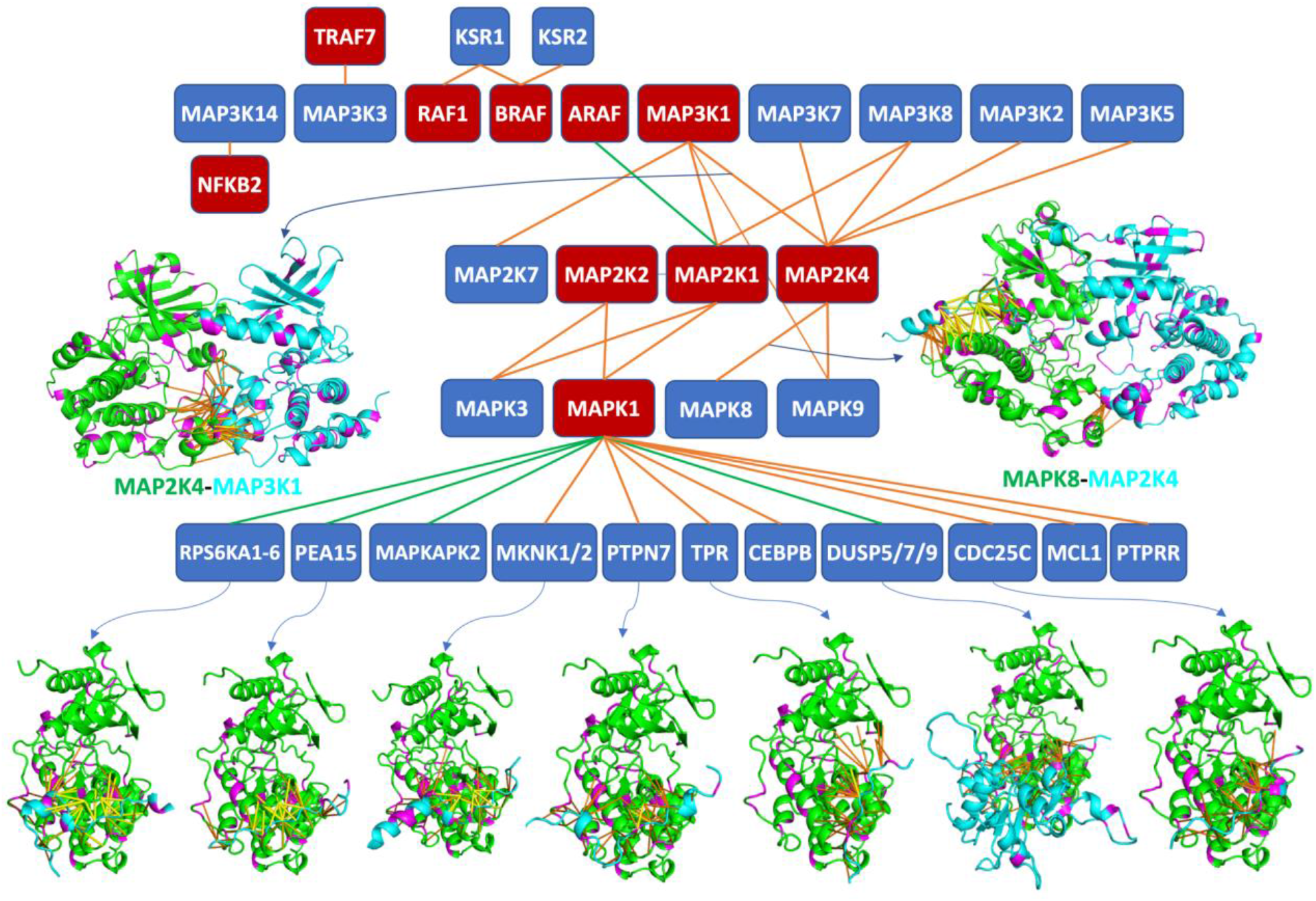
Interactions in the MAP kinase network captured by AlphaFold. This diagram shows the predicted interactions between members of the MAP kinase pathway. The MAP kinase cascades are arranged in four main layers: MAP3Ks (such as BRAF and MAP3K1), MAP2Ks (such as MAP2K1), MAPKs (such as MAPK1), and downstream targets and binding partners of MAPK1 (such as MKNK1). Cancer driving proteins are represented by red boxes, and their binding partners are represented by blue boxes. Lines connect protein pairs that are predicted to interact by AF, and pairs with and without known experimental structures are connected by green lines and orange lines, respectively. Structural models for some complexes, including MAP2K4-MAP3K1, MAPK8-MAP2K4, and MAPK1 (green) and some of its binding partners (cyan), are shown. In these models, predicted contacts are shown in sticks (yellow: probability score ≥ 0.9; orange: 0.5 ≤ probability score < 0.9). Positions with SMMs in cancers are shown in magenta.

Analysis of AF models suggests that interacting partners of MAPK1 mostly use a short linear sequence motif (docking site) to interact with a surface patch on the MAPK1 α-helical subdomain called the common docking (CD) domain [79] (Fig. 4). These docking sites, distal to the target phosphorylation sites, have been characterized in many MAPK substrates and other binding proteins [80-83]. One exception is that several phosphatases (DUS5/7/9) use a globular domain to interact with MAPK1 CD domain. The MAPK docking sites are also present in the N-terminal regions of upstream MAPK kinases (MAP2Ks or MEKs). In particular, the docking site in the N-terminal region of MAP2K4 shows high contact probabilities (>0.98) with MAP2K4’s substrates MAPK8 (shown in Fig. 4) and MAPK9.

The interfaces between MAPK1 and some of its binding partners (such as RPS6KA1, DUSP5 and MKNK2) are enriched with SMMs in cancer. This is partially due to the presence of a mutation hotspot residue E322 in the CD domain of MAPK1. Enrichment of SMMs at PPI interfaces were also observed for several pairs of MAP3Ks and MAP2Ks, such as MAP2K4-MAP3K1. MAP2K4 has a mutation hotspot residue R134 at its interface with MAP3K1 (Fig. 4). Oncogenic loss-of-function mutations in MAP2K4 and MAP3K1 have been identified in several types of cancers [21, 84, 85]. MAP3K1 has the dual function of pro-survival signaling and promotion of apoptosis [86]. MAP3K1 regulates the JNK pathway that can induce apoptosis to inhibit cancer growth through the MAP3K1-MAP2K4-JNK phosphorylation cascade, and inactivation of MAP3K1 and MAP2K4 was observed in certain cancers [86]. SMMs at the MAP3K1-MAP2K4 interface may disrupt the MAP3K1-MAP2K4-JNK signaling axis for apoptosis, antagonizing the tumor suppressor roles of MAP3K1 and MAP2K4.

### Structural insights into protein complexes in the Fanconi anemia pathway

DNA repair pathways play important roles in cancers. On the one hand, a number of genes functioning in DNA repair pathways have been classified as TSGs, such as BRCA1, BRCA2 [87], and genes causing the Bloom’s syndrome and the FA disease [88, 89]. Loss-of-function germline mutations or natural variants in these genes cause genomic instability and elevated rate of SMMs, resulting in an increased risk of developing various types of cancer. On the other hand, cancer cells utilize DNA repair pathways to help survive DNA damage induced by chemotherapeutic treatments. Therefore, inhibition of DNA repair pathways by targeting certain DNA repair genes could enhance the effects of chemotherapy [90, 91]. Since cells of certain tumors can only rely on a reduced set of DNA repair pathways for survival, drugs targeting these pathways could also be used as single-agent therapies for such cancer types [90, 91].

FA is a clinically heterogeneous genetic disorder characterized by congenital abnormalities, progressive bone marrow failure, chromosome instability, and an increased risk of developing tumors in early adulthood [89]. The FA pathway includes 22 FA proteins [92] whose mutations can lead to the FA disease (underlined proteins in Fig. 5). 12 FA genes have been classified as TSGs, including *FANCA, FANCC, FANCE, FANCF, FANCG, FANCD2, BRCA1* (*FANCS*), *BRCA2* (*FANCD1*), *BRIP1* (*FANCJ*), *PALB2* (*FANCN*), *ERCC4* (*FANCQ*), and *RFDW3* (*FANCW*) (red boxes in Fig. 5). FA proteins together with their interaction partners form protein complexes that are involved in the repair of DNA damages. AF identified a number of PPIs for proteins in the FA pathway, and many of them were not structurally characterized before (Fig. 5).

**Figure 5.**
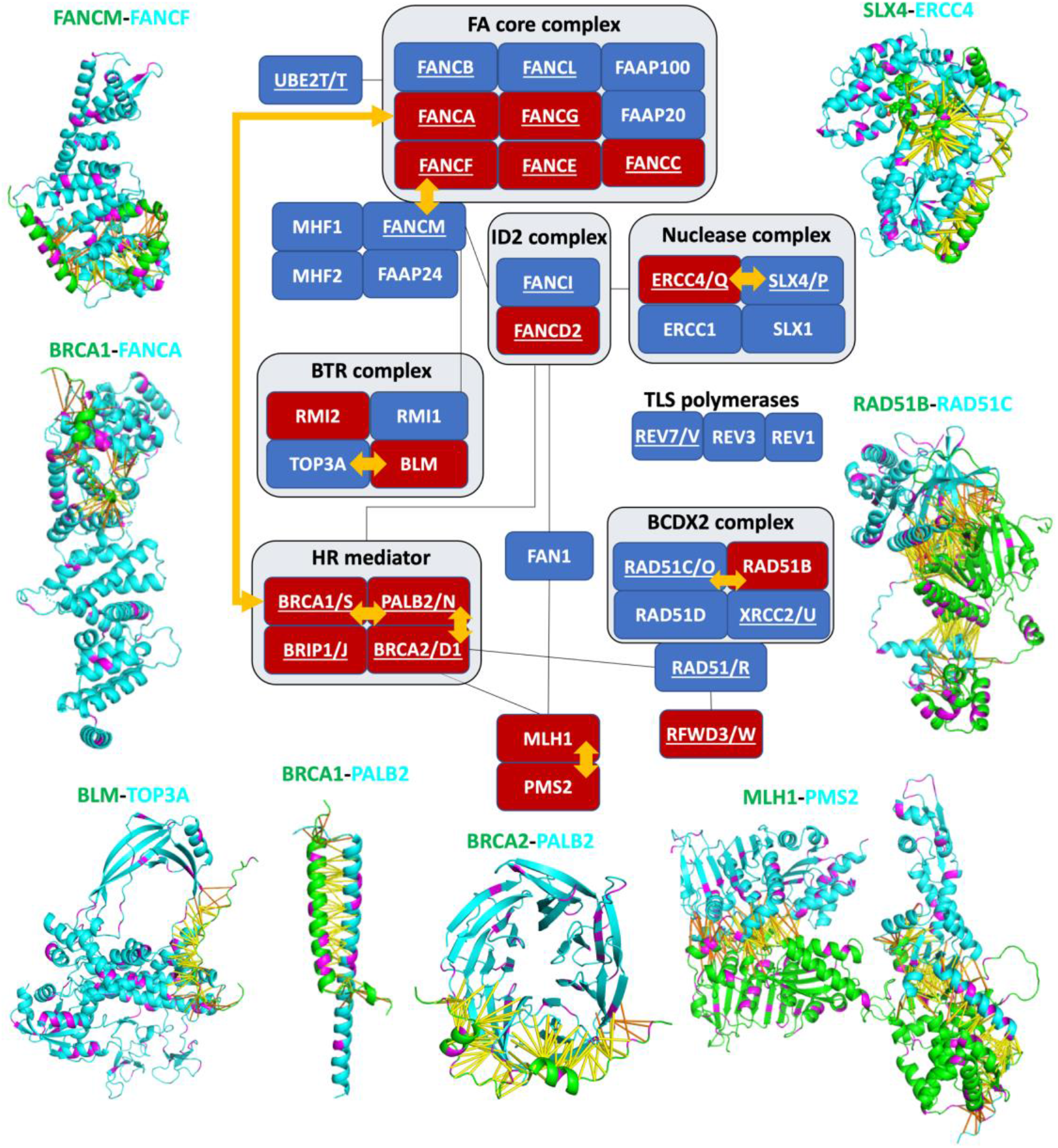
Structural insights into the proteins in the Fanconi anemia pathway. The FA pathway was adapted from KEGG and a review article [92]. Cancer drivers are represented as red boxes and other proteins are shown as blue boxes. Proteins forming complexes are arranged next to each other and additional interactions between components of different complexes are linked by black lines. FA proteins are underlined, and proteins classified as FA proteins but have other commonly used names are denoted by the FA name code after the slash (for example, BRCA1/S denote BRCA1 as FA proteins with name FANCS). AF predicted structural complexes without experimental structures are denoted by orange arrows between the interaction partners, and the AF structural models are shown for these pairs. In the shown structure models, positions with SMMs are colored in magenta. Predicted contacts are shown in sticks (yellow: probability score ≥ 0.9; orange: 0.5 ≤ probability score < 0.9).

The FA core complex is a large E3 ubiquitin ligase responsible for the monoubiquitylation of the FANCI-FANCD2 complex, which is a crucial step to recruit the latter to the DNA break site (Fig. 5). The structures of the FA core complexes from chicken and human were recently solved [93, 94]. FANCM forms a complex with FAAP24 and MHF1/2 to recruit the FA core complex to the DNA damage site. The interaction between FANCM and the FA core complex is through the FANCM-FANCF interaction. Despite previous efforts to map the interaction regions in FANCM and FANCF [95], the structure basis of this interaction is unknown. The AF model provides detailed information about the FANCM-FANCF interaction. Consistent with previous experimental results, the interface in FANCM is mapped to residues 960-1024, which forms three α-helices that wrap around the N-terminal α-helical domain of FANCF (residues 1-151) (Fig. 5).

SLX4 is a scaffold protein that together with ERCC1 and the two nucleases, SLX1 and ERCC4, forms a nuclease complex [96]. SLX4 (FANCP) and ERCC4 (FANCQ) have been classified as FA proteins as some of their mutations have the clinical features of the FA disease [97, 98]. AF predicted a highly confident model (contact probability > 0.99) of the SLX4-ERCC4 complex that lacks the experimental structure. In this model, SLX4 exploits several non-globular α-helices from residues 466-555 (green chain in Fig. 5) to interact with ERCC4, which is consistent with experimental studies that suggested that the interaction is mediated by the N-terminal region of SLX4, with residues 527-558 being critical [96, 99]. Conserved residues L530, F545, Y546 and L550 are directly involved in hydrophobic interactions in the interface in the AF model. Mutations in some of these positions such as L530Q and Y546C have been shown to enhance the degree of mitomycin C-induced chromosome aberrations [99].

BRCA2 (FANCD1) is a major player in the DNA repair pathways. BRCA2 forms a complex (Fig. 5) with BRCA1 (FANCS), PALB2 (FANCN) and BRIP1 (FANCJ) to mediate DNA repair through homologous recombination. A short N-terminal segment of BRCA2 (residues 24-37) has been shown to bind the beta-propeller domain of PALB2 by structural determination [100]. Mutations in this segment (G25R, W31R, W31C and F32L) were found to abolish the BRCA2-PALB2 interaction [101, 102]. The same segment mediates the BRCA2-PALB2 interaction in the AF model, where it forms a short α-helix in the middle that docks on the surface groove at one end of the fourth and fifth beta-propeller. Moreover, AF extended beyond the experimentally characterized BRCA2-PALB2 interface to include residues 7-23 of BRCA2, which interact with the surface groove at one end of the fifth and sixth beta-propeller of PALB2. The N-terminal interaction site of BRCA2 thus consists of a tandem repeat of a short helix and a loop behind it, which is reflected by the sequence similarity between the two repeats: both of them have the ϕϕExϕ motif (ϕ: a large hydrophobic residue, FFEIF and WFEEL in human BRCA2).

### Cancer drivers serve as hubs of protein-protein interactions

One prominent feature of the cancer interactome is the existence of proteins that interact with many other proteins, namely, PPI hubs (Table S40). Some of the PPI hubs, including the four largest hubs, EP300, BRCA1, CREBBP, and TRAPP, play central biological roles that require them to interact with many partners. However, some hubs stand out likely because the protein is prone to show false positive PPIs both in large-scale experiments and AF modeling. One example is EGFR, who had high AF contact probabilities with 32 proteins, but over half of them were mediated by its signal peptide that is expected to be cleaved. We hypothesized that the interactions mediated by this signal peptide represented false positives and removed them from our cancer interactome.

*BRCA1* is the most studied and most frequently mutated cancer-causing gene in hereditary breast cancer and ovarian cancer [103, 104]. BRCA1 has an N-terminal RING-type zinc finger domain (residues 1-103) with E3 ubiquitin ligase activity and two C-terminal BRCT domains (residues 1645-1863). These N- and C-terminal globular domains harbor most of the pathogenic mutations in BRCA1 [105]. The N-terminal RING domain of BRCA1 interacts with E2 ubiquitin-conjugating enzymes such as UBE2D3 and UBE2E2 [106] and other regulators such as BARD1 that enhances BRCA1’s E3 ligase activity [107, 108]. AF models of the RING domain and these binding partners (Fig. 6) resemble the experimentally determined complex structures [109, 110].

**Figure 6.**
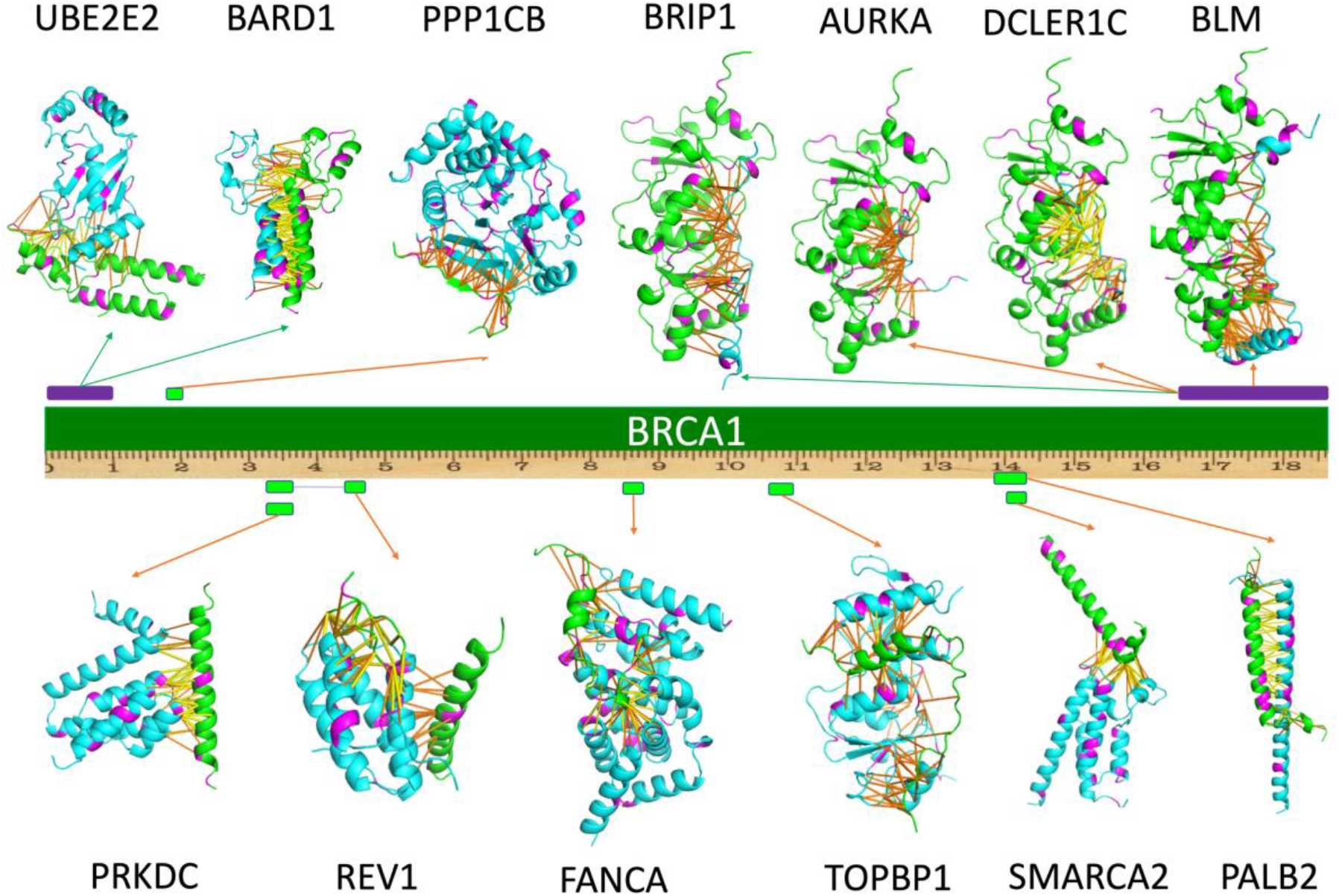
Examples of AlphaFold models of BRCA1 and its interacting partners. The BRCA1 protein is shown in the middle with a ruler to label the residue number (unit: 100 residues). The interaction regions in BRCA1 are shown as boxes above and below the BRCA1 protein. Purple boxes denote globular domains at the N- and C-terminal regions of BRCA1, and green boxes are from the middle region of BRCA1 without globular domains. Each arrow links a segment of BRCA1 to the structural model of the complex formed by this BRCA1 segment and its interaction partners. Green and orange arrows denote experimentally solved and unsolved structures, respectively. In all these AF models, BRCA1 is shown in green and its interaction partner is shown in cyan. Positions with SMMs are shown in purple. Predicted contacts are shown in sticks (yellow: probability score ≥ 0.9; orange: 0.5 ≤ probability score < 0.9).

Each C-terminal BRCT domain of BRCA1 consists of three α-helices surrounding four β-strands [111]. The relative positioning of the BRCT repeats allows them to recognize short sequence motifs containing phosphorylated Serine residues and aromatic residues (pSXXF motif) [112, 113]. AF models of BRCA1 BRCT domains with BRIP1, AURKA, and DCLER1C, respectively, closely mimic the experimentally determined complexes [113, 114], placing a Serine or Threonine residue (S990 of BRIP1, S51 of AURKA, and T557 of DCLER1C) to the location of phosphorylated Serine observed in experimental structures. Without knowing post-translational modifications, AF can correctly model such interactions mediated by phosphorylated peptides, possibly due to the coevolution between BRCT domains and residues in and around the pSXXF motifs. Interestingly, in the AF model, BLM does not utilize a pSXXF motif to interact with tandem BRCT domains, and it is questionable whether BLM has a pSXXF motif (Fig. S3). Instead, a loop from BLM occupies a location near the pSXXF-motif-binding site of the BRCT domains, and an α-helix to the N-terminus of this loop also contributes to the BRCA1-BLM interaction. Whether this alternative interacting mode is biologically relevant remains to be validated experimentally.

The majority of the middle region of BRCA1 is predicted to be disordered, but some segments have been found to interact with a number of key players in DNA repair, such as PALB2 and FANCA [115]. Experimental studies revealed that PALB2 interacts with BRCA1 using the N-terminal α-helix of PALB2 and residues 1385-1430 of BRCA1 [116, 117]. While this interaction was predicted to involve the formation of coiled coils, experimental structure of the BRCA1-PALB2 complex is absent. The AF model is consistent with this prediction and provides structural details of this coiled-coil interaction (Fig. 6). The BRCA1 segment interacting with PALB2 harbors three likely pathogenic mutations (Q1395H, Q1408H, and M1411H) classified in the ClinVar database [105], and PALB2 mutations in this interface such as L35P have been shown to compromise the BRCA1-PALB2 interaction and are associated with increased breast cancer risk [118].

The BRCA1-FANCA interaction has been experimentally mapped to the middle region of BRCA1 (740-1083) and the N-terminal region of FANCA (1-589) [119]. AF prediction is consistent with the experimental result, showing a conserved segment (848-877) of BRCA1 involved in the interaction with FANCA (Fig. 6). The FANCA interface also has a number of SMMs in cancer, including consecutive residues 420-422 (ESC) and 475-477 (TFL). The clinical significance of these mutations remains to be elucidated. AF also predicted several other interactions mapped to the middle region of BRCA1, involving non-globular sequence motifs in BRCA1 and globular domains found in proteins such as PRKDC, REV1, TOPBP1, and SMARCA2 (Fig. 6). Further characterization of these interactions may allow a deeper understanding of BRCA1 and its role in DNA repair and cancer.

## Materials and methods

### Identifying cancer-related protein-protein interactions from databases

The list of cancer driver proteins was obtained from the COSMIC cancer gene census database (https://cancer.sanger.ac.uk/census) (Oct 5th, 2021) [37]. We excluded cancer drivers whose “roles in cancer” were labeled solely as “fusion” by the COSMIC database, and mapped the remaining 588 proteins to 587 UniProt entries [120] (H3F3A and H3F3B mapped to the same entry). We obtained human PPIs from BIOGRID [8], IntAct [38], DIP [40], and MINT [39] databases, and we mapped proteins involved in these PPIs to 20,371 reviewed human entries in UniProt. These databases link each PPI to a set of PubMed IDs that represent the experimental studies supporting this interaction. We collected the PubMed IDs associated with each PPI from all databases. We counted the number of associated PPIs for each PubMed ID and considered a study to be a low-throughput if this number was no more than 10.

### Generation of protein sequence alignments

The eukaryotic proteomes were downloaded from the NCBI genome database. It consists of 49,102,568 proteins from 2,568 representative or reference proteomes (one proteome per species). We attempted to find orthologs from these proteomes for each reviewed human entry in the UniProt database [120]. For each human protein (referred to as the target protein below), the corresponding orthologous group at the eukaryotes level in OrthoDB [121] was identified. Human proteins that were not classified in OrthoDB or belonged to the Zinc finger C2H2-type group (OrthoDB group: 1318335at2759), the Reverse transcriptase, RNase H-like domain group (OrthoDB group: 583605at2759) and the Pentatricopeptide repeat group (Orthodb group: 1344243at2759) were ignored as these groups contain a large number (over 58000) of paralogous sequences from gene duplications. The set of OrthoDB orthologous proteins for any target protein were clustered by CD-HIT [122] at 40% identity level. For each CD-HIT cluster, we selected one representative sequence that showed the lowest BLAST [123] e-value to the target protein. The representative sequences were then used as queries to search against the eukaryotic proteomes to identify homologous proteins to the target protein (e-value cutoff: 0.001).

For each target protein, we identified a single best hit from each organism, if available, that satisfies two criteria: (1) its sequence identity to the target protein is above 35%; (2) it shows the highest sequence similarity to the target protein among the combined BLAST hits found by multiple representative sequences. These best hits together with the target protein were aligned by MAFFT (with the --auto option) [124]. We labeled each protein by the organism it belongs in the resulting multiple sequence alignments (MSAs) and removed positions that are gaps in the human proteins. To construct the joint MSA for any two human proteins, we concatenated the aligned sequences of the same organism from two MSAs. In cases where a sequence is missing from an organism, we used gaps to replace that sequence and added these cases at the end of the joint MSA.

### Modeling protein complexes with AlphaFold

One limitation of AF modeling is that large protein complexes cannot fit in the memory of our GPUs (≤ 48GB). To solve this problem, we parsed proteins into domains based on DeepMind’s models of monomeric human proteins (https://alphafold.ebi.ac.uk/). In addition to the predicted 3D structure, DeepMind also provided the predicted aligned error (PAE) for each residue pair in a protein. PAE reflects AF’s confidence in the distance between two residues, and it is suggested to be useful in defining domains of a protein [125]. Residues within the same domain are tightly packed together to form a globular and rigid 3D structure, and thus PAEs between them are expected to be low. In contrast, since the relative orientation and distance between different domains can be variable, a pair of residues from different domains frequently shows high PAE.

We wrote an in-house script (details in supplemental materials) to iterate the following procedure to split proteins into segments until all segments are shorter than 500 amino acids or cannot be further split. We split a protein or a segment into two segments if (1) it had more than 500 residues and (2) the density of residue pairs showing low PAE (≤12Å) within each segment (*D*_*intra*_) was much higher than the density of residue pairs showing low PAE (≤12Å) between two segments (*D*_*inte*r_). We found the split site that maximize the *D*_*intra*_ / *D*_*inter*_ ratio, and we required this ratio to be at least 10 for proteins or segments larger than 1000 residues and at least 20 for proteins or segments with 500-1000 residues.

As described previously, we deployed AF to model protein complexes by feeding it with joint MSAs of two proteins and introducing a gap of 200 residues in between. Protein pairs with combined length ≤ 1500 were directly modeled. Larger proteins were split into domains as described above and all pairs of domains were modeled if none of them has combined length > 1500. AF produces probabilities for the C_ß_ – C_ß_ distances of residue pairs to fall into a series of distance bins. Residue-residue contact probability was calculated as the sum of the probabilities for the distance bins under 12Å. For a pair of proteins, the matrix (*m*) of contact probability is of the shape (*L*_1_ + *L*_2_) by (*L*_1_ + *L*_2_), where *L*_1_ and *L*_2_ are the length of the first and second proteins, respectively. To investigate the inter protein contacts, we extracted the submatrix *m*^′^ = *m*[: *L*_1_][*L*_1_: *L*_1_ + *L*_2_] as shown in the red box of Fig. S4. The highest residue-residue contact probability in this matrix *m*′ is used as the contact probability for a pair of proteins.

### Performance evaluation

On the one hand, we sampled 8,000 pairs of candidate PPIs as positive control by two criteria: (1) at least one protein of each pair is a cancer driver, (2) the PPI is supported by at least one low-throughput study (associated with ≤ 10 PPIs in databases) or two high-throughput studies. On the other hand, we sampled 8,000 protein pairs as negative control, and each pair consisted of a cancer driver and a randomly selected protein that was not shown to interact with the cancer driver according to PPI databases. We applied AF to both the positive and negative control sets and compared the distribution of AF contact probabilities over these two sets.

To evaluate the accuracy of our 3D models of binary protein complexes, we used the protein complexes with closely related PDB templates. We considered a PDB entry to be a closely related template for a complex if both proteins are close homologs to different chains in this entry. We defined close homologs by the following criteria: (1) BLAST e-value < 0.00001, (2) sequence identity > 60%, and (3) query coverage > 50% or hit coverage > 50%. For each interacting residue pair predicted by AF with contact probability ≥ 0.2, we calculated the distance between their corresponding residues in the homologous templates. We used the minimal distance in templates as the observed distance for a residue pair. We considered a predicted contact to be correct if the observed distance is less than 12Å. The percentage of correctly predicted contacts in a AF model of protein complex was used as contact prediction accuracy for this model.

### Analysis of cancer interactome

We loaded 1798 interacting protein pairs with AF contact probability above 0.8 to the Cytoscape software [43], where proteins were represented as nodes and two nodes were connected by an edge if they interact. We used the community cluster method, Glay [45], to cluster the network formed by these PPIs assuming undirected edges. We performed functional enrichment analysis for clusters with ≥ 10 proteins using the “Database for Annotation, Visualization and Integrated Discovery” (DAVID) web server [126]. We annotated each cluster by the predominantly enriched functional group or the name of the protein involving in the largest number of PPIs within this cluster. In addition, we counted the number of SMMs in the pan-cancer genomic data for the cancer driver proteins in each cluster and tested whether they are enriched or depleted of SMMs using binomial tests (details below). Interesting biological examples from the cancer interactome were studied manually with the help of Pymol and literature.

### Mapping of somatic missense mutations in cancer on cancer interactome

We used the OncoVar database [36] that summarized the pan-cancer SMMs from The Cancer Genome Atlas (TCGA, 10,769 cases) and the International Cancer Genome Consortium (ICGC, 1,942 cases) projects. These SMMs were mapped to the human reference genome GRCh38. We mapped the 20,371 reviewed human UniProt entries to GRCh38 using genome annotation tracks provided by UniProt (https://www.uniprot.org/downloads) [120], and we focused on the 17,515 proteins that can be unambiguously mapped. These proteins contain 10,272,489 residue positions. A total of 1,901,488 SMMs were observed in these positions from the 12,711 cancer patients, indicating an average SMM rate of 0.00146% per position per patient.

Enrichment of SMMs in a position was assessed using a binomial test implemented by the scipy.stats module. The parameters given to the binomial test are: *X* = the total number of SMMs in this position, *n* = 12,711, i.e. total number of cancer patients, and *p* = 0.00146%, the average SMM rate. We corrected the statistical significance for multiple statistical tests by computing the false discovery rate (FDR) [127]. The corrected P-values (Q-values) are 0.0044 and 0.052 if there are 5 and 4 patients carrying mutations in this position, respectively, and we thus considered positions with ≥ 5 SMMs to be mutation hotspots. Similarly, we computed the enrichment of SMMs in sets of proteins using binomial tests, where *X* = the total number of mutations observed in these proteins, *n* = 12,711 x total number of positions in these proteins, and *p* = 0.00146%.

### Analysis of somatic missense mutation distribution in PPI interfaces

We considered AF-predicted contacts between proteins as residue pairs satisfying the following criteria: (1) the protein pair or domain pair that was modeled by AF has top contact probability ≥ 0.8; (2) this pair of residues has AF contact probability ≥ 0.2; (3) the distance between this residue pair in the AF 3D structure model is below 8Å. We considered residues that participated in any of the AF-predicted contacts as interface residues. We evaluated if an interface was enriched in or depleted of SMMs using a binomial test, where *X* = the total number of SMMs observed in the interface residues among all 12,711 patients, *n* = 12,711 x total number of interface residues, and *p* = 0.00146%. We corrected the statistical significance for multiple tests by computing FDR. We analyzed the functional preference for proteins contributing to the 177 interfaces that were enriched (P-value < 0.01, FDR < 0.1) in SMMs and proteins contributing to the 70 interfaces that were depleted (P-value < 0.01, FDR < 0.2) of SMMs using the DAVID web server. We used all proteins involved in our cancer interactome as the background for both functional enrichment analyses.

## References

1. Hanahan, D. and R.A. Weinberg, Hallmarks of cancer: the next generation. Cell, 2011. 144(5): p. 646–74.

2. Stelzl, U. and E.E. Wanker, The value of high quality protein-protein interaction networks for systems biology. Curr Opin Chem Biol, 2006. 10(6): p. 551–8.

3. Wells, J.A. and C.L. McClendon, Reaching for high-hanging fruit in drug discovery at protein-protein interfaces. Nature, 2007. 450(7172): p. 1001–9.

4. Ivanov, A.A., F.R. Khuri, and H. Fu, Targeting protein-protein interactions as an anticancer strategy. Trends Pharmacol Sci, 2013. 34(7): p. 393–400.

5. Lu, H., et al., Recent advances in the development of protein-protein interactions modulators: mechanisms and clinical trials. Signal Transduct Target Ther, 2020. 5(1): p. 213.

6. Rolland, T., et al., A proteome-scale map of the human interactome network. Cell, 2014. 159(5): p. 1212–1226.

7. Rual, J.F., et al., Towards a proteome-scale map of the human protein-protein interaction network. Nature, 2005. 437(7062): p. 1173–8.

8. Oughtred, R., et al., The BioGRID interaction database: 2019 update. Nucleic Acids Res, 2019. 47(D1): p. D529–D541.

9. Li, Z., et al., The OncoPPi network of cancer-focused protein-protein interactions to inform biological insights and therapeutic strategies. Nat Commun, 2017. 8: p. 14356.

10. Kim, M., et al., A protein interaction landscape of breast cancer. Science, 2021. 374(6563): p. eabf3066.

11. Kuchaiev, O., et al., Geometric de-noising of protein-protein interaction networks. PLoS Comput Biol, 2009. 5(8): p. e1000454.

12. Mackay, J.P., et al., Protein interactions: is seeing believing? Trends Biochem Sci, 2007. 32(12): p. 530–1.

13. Chin, L., et al., Making sense of cancer genomic data. Genes Dev, 2011. 25(6): p. 534–55.

14. Vogelstein, B., et al., Cancer genome landscapes. Science, 2013. 339(6127): p. 1546–58.

15. Bailey, M.H., et al., Comprehensive Characterization of Cancer Driver Genes and Mutations. Cell, 2018. 173(2): p. 371–385 e18.

16. Tate, J.G., et al., COSMIC: the Catalogue Of Somatic Mutations In Cancer. Nucleic Acids Res, 2019. 47(D1): p. D941–D947.

17. International Cancer Genome, C., et al., International network of cancer genome projects. Nature, 2010. 464(7291): p. 993–8.

18. Cancer Genome Atlas Research, N., et al., The Cancer Genome Atlas Pan-Cancer analysis project. Nat Genet, 2013. 45(10): p. 1113–20.

19. Nakagawa, H., et al., Cancer whole-genome sequencing: present and future. Oncogene, 2015. 34(49): p. 5943–50.

20. Porta-Pardo, E., A. Valencia, and A. Godzik, Understanding oncogenicity of cancer driver genes and mutations in the cancer genomics era. FEBS Lett, 2020. 594(24): p. 4233–4246.

21. Pereira, B., et al., The somatic mutation profiles of 2,433 breast cancers refines their genomic and transcriptomic landscapes. Nat Commun, 2016. 7: p. 11479.

22. Brennan, C.W., et al., The somatic genomic landscape of glioblastoma. Cell, 2013. 155(2): p. 462–77.

23. Chang, M.T., et al., Identifying recurrent mutations in cancer reveals widespread lineage diversity and mutational specificity. Nat Biotechnol, 2016. 34(2): p. 155–63.

24. Engin, H.B., J.F. Kreisberg, and H. Carter, Structure-Based Analysis Reveals Cancer Missense Mutations Target Protein Interaction Interfaces. PLoS One, 2016. 11(4): p. e0152929.

25. Gao, J., et al., 3D clusters of somatic mutations in cancer reveal numerous rare mutations as functional targets. Genome Med, 2017. 9(1): p. 4.

26. Porta-Pardo, E., et al., A Pan-Cancer Catalogue of Cancer Driver Protein Interaction Interfaces. PLoS Comput Biol, 2015. 11(10): p. e1004518.

27. Nishi, H., et al., Cancer missense mutations alter binding properties of proteins and their interaction networks. PLoS One, 2013. 8(6): p. e66273.

28. Weis, K.E., et al., Constitutively active human estrogen receptors containing amino acid substitutions for tyrosine 537 in the receptor protein. Mol Endocrinol, 1996. 10(11): p. 1388–98.

29. Littlefield, P., et al., Structural analysis of the EGFR/HER3 heterodimer reveals the molecular basis for activating HER3 mutations. Sci Signal, 2014. 7(354): p. ra114.

30. Pahuja, K.B., et al., Actionable Activating Oncogenic ERBB2/HER2 Transmembrane and Juxtamembrane Domain Mutations. Cancer Cell, 2018. 34(5): p. 792–806 e5.

31. Tunyasuvunakool, K., et al., Highly accurate protein structure prediction for the human proteome. Nature, 2021. 596(7873): p. 590–596.

32. Burke, D., et al., Towards a structurally resolved human protein interaction network. bioRxiv, 2021. DOI: 10.1101/2021.11.08.467664

33. Baek, M., et al., Accurate prediction of protein structures and interactions using a three-track neural network. Science, 2021. 373(6557): p. 871–876.

34. Jumper, J., et al., Highly accurate protein structure prediction with AlphaFold. Nature, 2021. 596(7873): p. 583–589.

35. Humphreys, I.R., et al., Computed structures of core eukaryotic protein complexes. Science, 2021. 374(6573): p. eabm4805.

36. Wang, T., et al., OncoVar: an integrated database and analysis platform for oncogenic driver variants in cancers. Nucleic Acids Res, 2021. 49(D1): p. D1289–D1301.

37. Sondka, Z., et al., The COSMIC Cancer Gene Census: describing genetic dysfunction across all human cancers. Nat Rev Cancer, 2018. 18(11): p. 696–705.

38. Hermjakob, H., et al., IntAct: an open source molecular interaction database. Nucleic Acids Res, 2004. 32(Database issue): p. D452–5.

39. Zanzoni, A., et al., MINT: a Molecular INTeraction database. FEBS Lett, 2002. 513(1): p. 135–40.

40. Xenarios, I., et al., DIP: the database of interacting proteins. Nucleic Acids Res, 2000. 28(1): p. 289–91.

41. Sun, J. and Z. Zhao, A comparative study of cancer proteins in the human protein-protein interaction network. BMC Genomics, 2010. 11 Suppl 3: p. S5.

42. Jonsson, P.F. and P.A. Bates, Global topological features of cancer proteins in the human interactome. Bioinformatics, 2006. 22(18): p. 2291–7.

43. Shannon, P., et al., Cytoscape: a software environment for integrated models of biomolecular interaction networks. Genome Res, 2003. 13(11): p. 2498–504.

44. Du, Y., et al., PINA 3.0: mining cancer interactome. Nucleic Acids Res, 2021. 49(D1): p. D1351–D1357.

45. Su, G., et al., GLay: community structure analysis of biological networks. Bioinformatics, 2010. 26(24): p. 3135–7.

46. Nair, S.S. and R. Kumar, Chromatin remodeling in cancer: a gateway to regulate gene transcription. Mol Oncol, 2012. 6(6): p. 611–9.

47. Lancaster, J.M., et al., BRCA2 mutations in primary breast and ovarian cancers. Nat Genet, 1996. 13(2): p. 238–40.

48. Frank, T.S., et al., Clinical characteristics of individuals with germline mutations in BRCA1 and BRCA2: analysis of 10,000 individuals. J Clin Oncol, 2002. 20(6): p. 1480–90.

49. Hosoya, N. and K. Miyagawa, Targeting DNA damage response in cancer therapy. Cancer Sci, 2014. 105(4): p. 370–88.

50. Kamburov, A., et al., Comprehensive assessment of cancer missense mutation clustering in protein structures. Proc Natl Acad Sci U S A, 2015. 112(40): p. E5486–95.

51. Olingy, C.E., H.Q. Dinh, and C.C. Hedrick, Monocyte heterogeneity and functions in cancer. J Leukoc Biol, 2019. 106(2): p. 309–322.

52. Yuen, G.J., E. Demissie, and S. Pillai, B lymphocytes and cancer: a love-hate relationship. Trends Cancer, 2016. 2(12): p. 747–757.

53. Serrano, M., The tumor suppressor protein p16INK4a. Exp Cell Res, 1997. 237(1): p. 7–13.

54. Ortega, S., M. Malumbres, and M. Barbacid, Cyclin D-dependent kinases, INK4 inhibitors and cancer. Biochim Biophys Acta, 2002. 1602(1): p. 73–87.

55. Jeffrey, P.D., L. Tong, and N.P. Pavletich, Structural basis of inhibition of CDK-cyclin complexes by INK4 inhibitors. Genes Dev, 2000. 14(24): p. 3115–25.

56. Russo, A.A., et al., Structural basis for inhibition of the cyclin-dependent kinase Cdk6 by the tumour suppressor p16INK4a. Nature, 1998. 395(6699): p. 237–43.

57. Shiloh, Y. and Y. Ziv, The ATM protein kinase: regulating the cellular response to genotoxic stress, and more. Nat Rev Mol Cell Biol, 2013. 14(4): p. 197–210.

58. Wardlaw, C.P., A.M. Carr, and A.W. Oliver, TopBP1: A BRCT-scaffold protein functioning in multiple cellular pathways. DNA Repair (Amst), 2014. 22: p. 165–74.

59. Stankovic, T., et al., ATM mutations and phenotypes in ataxia-telangiectasia families in the British Isles: expression of mutant ATM and the risk of leukemia, lymphoma, and breast cancer. Am J Hum Genet, 1998. 62(2): p. 334–45.

60. Becker-Catania, S.G., et al., Ataxia-telangiectasia: phenotype/genotype studies of ATM protein expression, mutations, and radiosensitivity. Mol Genet Metab, 2000. 70(2): p. 122–33.

61. Li, Y., et al., UTRN on chromosome 6q24 is mutated in multiple tumors. Oncogene, 2007. 26(42): p. 6220–8.

62. Vizkeleti, L., et al., Altered integrin expression patterns shown by microarray in human cutaneous melanoma. Melanoma Res, 2017. 27(3): p. 180–188.

63. Zhou, S., et al., UTRN inhibits melanoma growth by suppressing p38 and JNK/c-Jun signaling pathways. Cancer Cell Int, 2021. 21(1): p. 88.

64. Oster, S.K., et al., The myc oncogene: MarvelouslY Complex. Adv Cancer Res, 2002. 84: p. 81–154.

65. Burnichon, N., et al., MAX mutations cause hereditary and sporadic pheochromocytoma and paraganglioma. Clin Cancer Res, 2012. 18(10): p. 2828–37.

66. Comino-Mendez, I., et al., Functional and in silico assessment of MAX variants of unknown significance. J Mol Med (Berl), 2015. 93(11): p. 1247–55.

67. Romero, O.A., et al., MAX inactivation in small cell lung cancer disrupts MYC-SWI/SNF programs and is synthetic lethal with BRG1. Cancer Discov, 2014. 4(3): p. 292–303.

68. Augert, A., et al., MAX Functions as a Tumor Suppressor and Rewires Metabolism in Small Cell Lung Cancer. Cancer Cell, 2020. 38(1): p. 97–114 e7.

69. Denslow, S.A. and P.A. Wade, The human Mi-2/NuRD complex and gene regulation. Oncogene, 2007. 26(37): p. 5433–8.

70. Xia, L., et al., CHD4 Has Oncogenic Functions in Initiating and Maintaining Epigenetic Suppression of Multiple Tumor Suppressor Genes. Cancer Cell, 2017. 31(5): p. 653–668 e7.

71. Spruijt, C.G., et al., Cross-linking mass spectrometry reveals the structural topology of peripheral NuRD subunits relative to the core complex. FEBS J, 2021. 288(10): p. 3231–3245.

72. Roy, H.K., et al., AKT proto-oncogene overexpression is an early event during sporadic colon carcinogenesis. Carcinogenesis, 2002. 23(1): p. 201–5.

73. Altomare, D.A. and J.R. Testa, Perturbations of the AKT signaling pathway in human cancer. Oncogene, 2005. 24(50): p. 7455–64.

74. Fu, Z. and D.J. Tindall, FOXOs, cancer and regulation of apoptosis. Oncogene, 2008. 27(16): p. 2312–9.

75. Calnan, D.R. and A. Brunet, The FoxO code. Oncogene, 2008. 27(16): p. 2276–88.

76. Avruch, J., MAP kinase pathways: the first twenty years. Biochim Biophys Acta, 2007. 1773(8): p. 1150–60.

77. Davies, H., et al., Mutations of the BRAF gene in human cancer. Nature, 2002. 417(6892): p. 949–54.

78. Prior, I.A., P.D. Lewis, and C. Mattos, A comprehensive survey of Ras mutations in cancer. Cancer Res, 2012. 72(10): p. 2457–67.

79. Arvind, R., et al., A mutation in the common docking domain of ERK2 in a human cancer cell line, which was associated with its constitutive phosphorylation. Int J Oncol, 2005. 27(6): p. 1499–504.

80. Jacobs, D., et al., Multiple docking sites on substrate proteins form a modular system that mediates recognition by ERK MAP kinase. Genes Dev, 1999. 13(2): p. 163–75.

81. Akella, R., T.M. Moon, and E.J. Goldsmith, Unique MAP Kinase binding sites. Biochim Biophys Acta, 2008. 1784(1): p. 48–55.

82. Sharrocks, A.D., S.H. Yang, and A. Galanis, Docking domains and substrate-specificity determination for MAP kinases. Trends Biochem Sci, 2000. 25(9): p. 448–53.

83. Bardwell, L., Mechanisms of MAPK signalling specificity. Biochem Soc Trans, 2006. 34(Pt 5): p. 837–41.

84. Michaut, M., et al., Integration of genomic, transcriptomic and proteomic data identifies two biologically distinct subtypes of invasive lobular breast cancer. Sci Rep, 2016. 6: p. 18517.

85. Ciriello, G., et al., Comprehensive Molecular Portraits of Invasive Lobular Breast Cancer. Cell, 2015. 163(2): p. 506–19.

86. Pham, T.T., S.P. Angus, and G.L. Johnson, MAP3K1: Genomic Alterations in Cancer and Function in Promoting Cell Survival or Apoptosis. Genes Cancer, 2013. 4(11-12): p. 419–26.

87. Venkitaraman, A.R., Cancer susceptibility and the functions of BRCA1 and BRCA2. Cell, 2002. 108(2): p. 171–82.

88. Ellis, N.A. and J. German, Molecular genetics of Bloom’s syndrome. Hum Mol Genet, 1996. 5 Spec No: p. 1457–63.

89. Ceccaldi, R., P. Sarangi, and A.D. D’Andrea, The Fanconi anaemia pathway: new players and new functions. Nat Rev Mol Cell Biol, 2016. 17(6): p. 337–49.

90. Helleday, T., et al., DNA repair pathways as targets for cancer therapy. Nat Rev Cancer, 2008. 8(3): p. 193–204.

91. Curtin, N.J., DNA repair dysregulation from cancer driver to therapeutic target. Nat Rev Cancer, 2012. 12(12): p. 801–17.

92. Rageul, J. and H. Kim, Fanconi anemia and the underlying causes of genomic instability. Environ Mol Mutagen, 2020. 61(7): p. 693–708.

93. Shakeel, S., et al., Structure of the Fanconi anaemia monoubiquitin ligase complex. Nature, 2019. 575(7781): p. 234–237.

94. Wang, S., et al., Structure of the FA core ubiquitin ligase closing the ID clamp on DNA. Nat Struct Mol Biol, 2021. 28(3): p. 300–309.

95. Deans, A.J. and S.C. West, FANCM connects the genome instability disorders Bloom’s Syndrome and Fanconi Anemia. Mol Cell, 2009. 36(6): p. 943–53.

96. Fekairi, S., et al., Human SLX4 is a Holliday junction resolvase subunit that binds multiple DNA repair/recombination endonucleases. Cell, 2009. 138(1): p. 78–89.

97. Bogliolo, M., et al., Mutations in ERCC4, encoding the DNA-repair endonuclease XPF, cause Fanconi anemia. Am J Hum Genet, 2013. 92(5): p. 800–6.

98. Kim, Y., et al., Mutations of the SLX4 gene in Fanconi anemia. Nat Genet, 2011. 43(2): p. 142–6.

99. Hashimoto, K., et al., Physical interaction between SLX4 (FANCP) and XPF (FANCQ) proteins and biological consequences of interaction-defective missense mutations. DNA Repair (Amst), 2015. 35: p. 48–54.

100. Oliver, A.W., et al., Structural basis for recruitment of BRCA2 by PALB2. EMBO Rep, 2009. 10(9): p. 990–6.

101. Xia, B., et al., Control of BRCA2 cellular and clinical functions by a nuclear partner, PALB2. Mol Cell, 2006. 22(6): p. 719–729.

102. Katagiri, T., et al., High proportion of missense mutations of the BRCA1 and BRCA2 genes in Japanese breast cancer families. J Hum Genet, 1998. 43(1): p. 42–8.

103. Narod, S.A. and W.D. Foulkes, BRCA1 and BRCA2: 1994 and beyond. Nat Rev Cancer, 2004. 4(9): p. 665–76.

104. Takaoka, M. and Y. Miki, BRCA1 gene: function and deficiency. Int J Clin Oncol, 2018. 23(1): p. 36–44.

105. Pei, J. and N.V. Grishin, The DBSAV Database: Predicting Deleteriousness of Single Amino Acid Variations in the Human Proteome. J Mol Biol, 2021. 433(11): p. 166915.

106. Wu, W., et al., The ubiquitin E3 ligase activity of BRCA1 and its biological functions. Cell Div, 2008. 3: p. 1.

107. Wu, L.C., et al., Identification of a RING protein that can interact in vivo with the BRCA1 gene product. Nat Genet, 1996. 14(4): p. 430–40.

108. Xia, Y., et al., Enhancement of BRCA1 E3 ubiquitin ligase activity through direct interaction with the BARD1 protein. J Biol Chem, 2003. 278(7): p. 5255–63.

109. Hu, Q., et al., Mechanisms of BRCA1-BARD1 nucleosome recognition and ubiquitylation. Nature, 2021. 596(7872): p. 438–443.

110. Brzovic, P.S., et al., Structure of a BRCA1-BARD1 heterodimeric RING-RING complex. Nat Struct Biol, 2001. 8(10): p. 833–7.

111. Williams, R.S., R. Green, and J.N. Glover, Crystal structure of the BRCT repeat region from the breast cancer-associated protein BRCA1. Nat Struct Biol, 2001. 8(10): p. 838–42.

112. Williams, R.S., et al., Structural basis of phosphopeptide recognition by the BRCT domain of BRCA1. Nat Struct Mol Biol, 2004. 11(6): p. 519–25.

113. Clapperton, J.A., et al., Structure and mechanism of BRCA1 BRCT domain recognition of phosphorylated BACH1 with implications for cancer. Nat Struct Mol Biol, 2004. 11(6): p. 512–8.

114. Varma, A.K., et al., Structural basis for cell cycle checkpoint control by the BRCA1-CtIP complex. Biochemistry, 2005. 44(33): p. 10941–6.

115. Clark, S.L., et al., Structure-Function Of The Tumor Suppressor BRCA1. Comput Struct Biotechnol J, 2012. 1(1).

116. Zhang, F., et al., PALB2 links BRCA1 and BRCA2 in the DNA-damage response. Curr Biol, 2009. 19(6): p. 524–9.

117. Sy, S.M., M.S. Huen, and J. Chen, PALB2 is an integral component of the BRCA complex required for homologous recombination repair. Proc Natl Acad Sci U S A, 2009. 106(17): p. 7155–60.

118. Foo, T.K., et al., Compromised BRCA1-PALB2 interaction is associated with breast cancer risk. Oncogene, 2017. 36(29): p. 4161–4170.

119. Folias, A., et al., BRCA1 interacts directly with the Fanconi anemia protein FANCA. Hum Mol Genet, 2002. 11(21): p. 2591–7.

120. UniProt, C., UniProt: the universal protein knowledgebase in 2021. Nucleic Acids Res, 2021. 49(D1): p. D480–D489.

121. Kriventseva, E.V., et al., OrthoDB v10: sampling the diversity of animal, plant, fungal, protist, bacterial and viral genomes for evolutionary and functional annotations of orthologs. Nucleic Acids Res, 2019. 47(D1): p. D807–D811.

122. Fu, L., et al., CD-HIT: accelerated for clustering the next-generation sequencing data. Bioinformatics, 2012. 28(23): p. 3150–2.

123. Altschul, S.F., et al., Gapped BLAST and PSI-BLAST: a new generation of protein database search programs. Nucleic Acids Res, 1997. 25(17): p. 3389–402.

124. Katoh, K. and D.M. Standley, MAFFT multiple sequence alignment software version 7: improvements in performance and usability. Mol Biol Evol, 2013. 30(4): p. 772–80.

125. Varadi, M., et al., AlphaFold Protein Structure Database: massively expanding the structural coverage of protein-sequence space with high-accuracy models. Nucleic Acids Res, 2021.

126. Huang da, W., B.T. Sherman, and R.A. Lempicki, Systematic and integrative analysis of large gene lists using DAVID bioinformatics resources. Nat Protoc, 2009. 4(1): p. 44–57.

127. Benjamini, Y. and Y. Hochberg, Controlling the false discovery rate: A practical and powerful approach to multiple testing. Journal of the Royal Statistical Society. Series B (Methodological), 1995. 57(1): p. 289–300.

